# Genome-wide meQTL mapping in cattle blood reveals *cis* and *trans* regulation of DNA methylation

**DOI:** 10.64898/2026.07.07.736355

**Authors:** Corentin Fouéré, Valentin Costes, Florian Besnard, Chrystelle Le Danvic, Clotilde Patry, Sébastien Fritz, Mekki Boussaha, Mélanie Jouin, Didier Boichard, Hélène Kiefer, Gabriel Costa Monteiro Moreira, Marie-Pierre Sanchez

## Abstract

**Background:** Complex traits are influenced by numerous variants, most of which have regulatory effects on gene expression that can be mediated by DNA methylation. Molecular QTL mapping is an approach that aims to dissect these effects. However, obtaining molecular phenotypes on a large scale is challenging, particularly in livestock species. In cattle, an epigenotyping array called EpiChip has recently been developed in the European RUMIGEN project. The EpiChip, which contains 43,317 CpG sites distributed all over the bovine genome, enables large-scale measurement of DNA methylation. This study aims to characterize the genetic determinism of blood DNA methylation in cows by estimating heritability and mapping *cis-* and *trans-*methylation QTLs (meQTLs).

**Results:** Whole blood samples from 4,457 genotyped Holstein cows were epigenotyped. Across all CpG sites, the heritability estimates averaged 24.6%. The local meQTL mapping at sequence-level for variable CpG sites (SD > 2.5%; n = 28,806) detected *cis*-meQTLs for 80.1% of the CpG sites, with sentinel SNPs located close to their associated CpGs. A two-step analysis was also conducted to identify long-range associations, with a particular focus on *trans*-meQTL hotspots. First, we identified CpG-SNP *trans*-associations using medium-density genotypes (50k SNPs) that revealed 31,846 SNPs with significant effects on 1 to 530 *trans-*CpG sites. Then, regions associated with at least 34 independent *trans*-CpGs were retained defining 31 hotpots. For each hotspot, a local sequence-level GWAS was conducted using the first principal component derived from the associated *trans*-CpGs. Out of the 31 detected hotspots, three were located close to transcription factor genes (*RUNX1*, *NFIC* and *FOXA3*) for which the associated *trans*-CpGs were enriched for the corresponding binding motif. Two other hotspots were located within *KDM5A* and *KDM5B*, and their corresponding *trans*-CpGs were strongly overrepresented in H3K4me3 narrow peaks in blood as well as in other tissues.

**Conclusions:** By identifying functional candidate genes associated with blood DNA methylation in cattle, these findings provide new insights into the regulatory architecture of DNA methylation in mammals, highlighting the value of large-scale molecular data from livestock populations.

## Introduction

Cattle is a livestock species of major economic importance and a powerful model for studying the genetic architecture of complex traits in mammals. The availability of large phenotypic and genomics datasets, often comprising tens to hundreds of thousands of individuals, has enabled extensive genome-wide association analyses that have identified numerous quantitative trait loci (QTL) affecting a wide range of traits of interest. Although several causal variants have been mapped to coding regions, it is now widely recognized that a substantial proportion of the genetic variation underlying complex traits is mediated by regulatory variants that influence gene expression and other molecular processes [1–3]. However, identifying these regulatory variants and elucidating the molecular mechanisms through which they affect phenotypes remain challenging. To address this gap, large-scale initiatives have been undertaken to generate functional genomic resources in cattle [4–6], with the aim of linking regulatory variation to intermediate molecular phenotypes and ultimately complex traits. Such resources facilitate the characterization of molecular QTL (molQTL) and provide novel insights into the regulatory architecture of the genome [6–8].

Among the molQTLs, expression QTLs (eQTLs) and splicing QTLs (sQTLs) have been the most extensively studied in cattle, revealing the wide genetic control of transcriptional regulation [7,9]. In contrast, the genetic determinants of DNA methylation variation remain much less explored, despite the key role of DNA methylation as a major epigenetic mechanism involved in gene regulation, chromatin organization, and genome stability. Mapping methylation QTLs (meQTL) provides a powerful framework to investigate how genetic variants influence the epigenome and to uncover regulatory pathways linking genetic variation to phenotypic diversity. In humans, large-scale studies have demonstrated that meQTLs are abundant across the genome and have provided valuable insights into the genetic architecture of complex traits and diseases [10–14]. However, similar investigations in livestock species have been limited, largely due to the cost and technical challenges associated with measuring DNA methylation in large numbers of animals. Recently, the development of the RUMIGEN EpiChip has opened new opportunities for large-scale epigenomic studies in cattle. This chip targets 43,317 CpG sites distributed across the genome and was designed on data from different studies comprising CpGs where methylation varies according to health status, physiological stage, fertility and environmental challenges, as well as CpGs co-localized with regulatory elements [15].

In this study, we leveraged a cohort of 4,457 cows epigenotyped using the RUMIGEN EpiChip and genotyped with a medium-density BeadChip in order to investigate the genetic regulation of DNA methylation in blood. By combining meQTL mapping with the integration of functional genomic information, we identified several candidate regulators with genome-wide effects, highlighting different regulatory mechanisms that contribute to epigenetic variation in cattle.

## Results

We investigated the genetic regulation of blood DNA methylation by estimating the variance components of methylation rates and identifying meQTLs. The main steps are presented in Fig. 1. For each CpG site targeted by the EpiChip, heritability was estimated using a genomic relationship matrix. Methylation QTL mapping was performed for the probes that display inter-individual variations (SD>2.5%), corresponding to 28,806 CpG sites. *Cis*-meQTL mapping was done at sequence level from imputed medium-density genotypes. *Trans*-meQTL mapping was done at the medium density. Genomic regions associated with numerous *trans*-CpGs were then re-assessed at the sequence level. In this second analysis, the coordinates of the cows along the first principal component of the *trans*-associated CpGs were used as a summarizing variable (see “Methods” section for further details).

**Figure 1.**
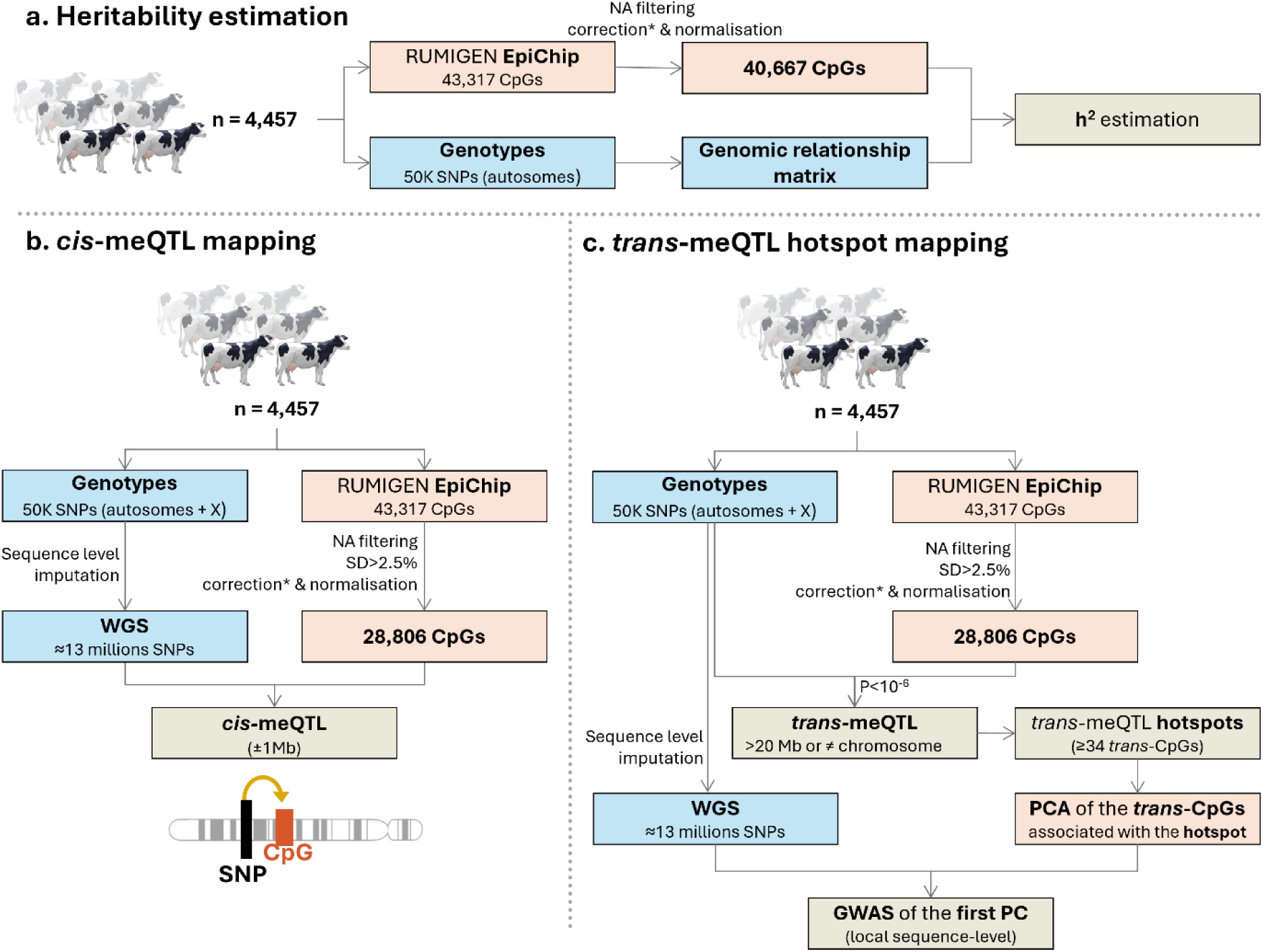
Study design. (a) The heritability of the CpG sites targeted by the EpiChip was estimated using the genomic relationship matrix. (b-c) Variable CpGs (SD > 2.5%) were investigated for (b) *cis*-meQTLs (sequence-level GWAS within ±1 Mb of the CpG) or (c) *trans*-meQTLs (medium-density GWAS). A second step was done for the *trans*-meQTL mapping process, focusing on regions where *trans*-meQTLs showed an association with at least 34 independent (>500 kbp apart) *trans*-CpGs (*trans*-meQTL hotspots). For each hotspot, we performed a Principal Component Analysis (PCA) of the *trans*-CpGs and considered the CpG coordinates on PC1 for a sequence-level GWAS of a 4 Mb region centered on the sentinel SNP (SNP associated with the highest number of *trans*-CpGs). correction*: cow age, blood formula, position of the chip on the slide and treatment batch.

### Heritability of DNA methylation at the CpG sites targeted by the EpiChip

Among the 40,676 CpG sites analyzed, the average heritability was 24.6% (SD = 22.5%), with 64.5% of CpGs showing a heritability greater than 10%, indicating that a substantial proportion of DNA methylation variation is under genetic control. However, heritability estimates were highly variables across CpG sites, ranging from zero to nearly one (Fig. 2a). CpGs annotated within promoter regions displayed low heritability estimates while those located in enhancer regions tended to have higher heritability estimates (Fig. 2c). In contrast, differences in heritability distributions across the CpG island landscape were less pronounced (Fig. 2b). Chang et al. [16] identified correlated regions of systemic interindividual epigenetic variation (CoRSIVs) in cattle, strongly associated with genetic variation, as observed for humans CoRSIVs. The 19 CpGs located within CoRSIVs showed a mean heritability of 43.1% (SD = 23.6%), which was significantly higher that observed for CpGs outside CoRSIVs (*P* = 5.3 × 10^-3^, two-sample t-test). Detailed heritability estimates for all CpG sites are provided in Additional file 1: Table S1.

**Figure 2.**
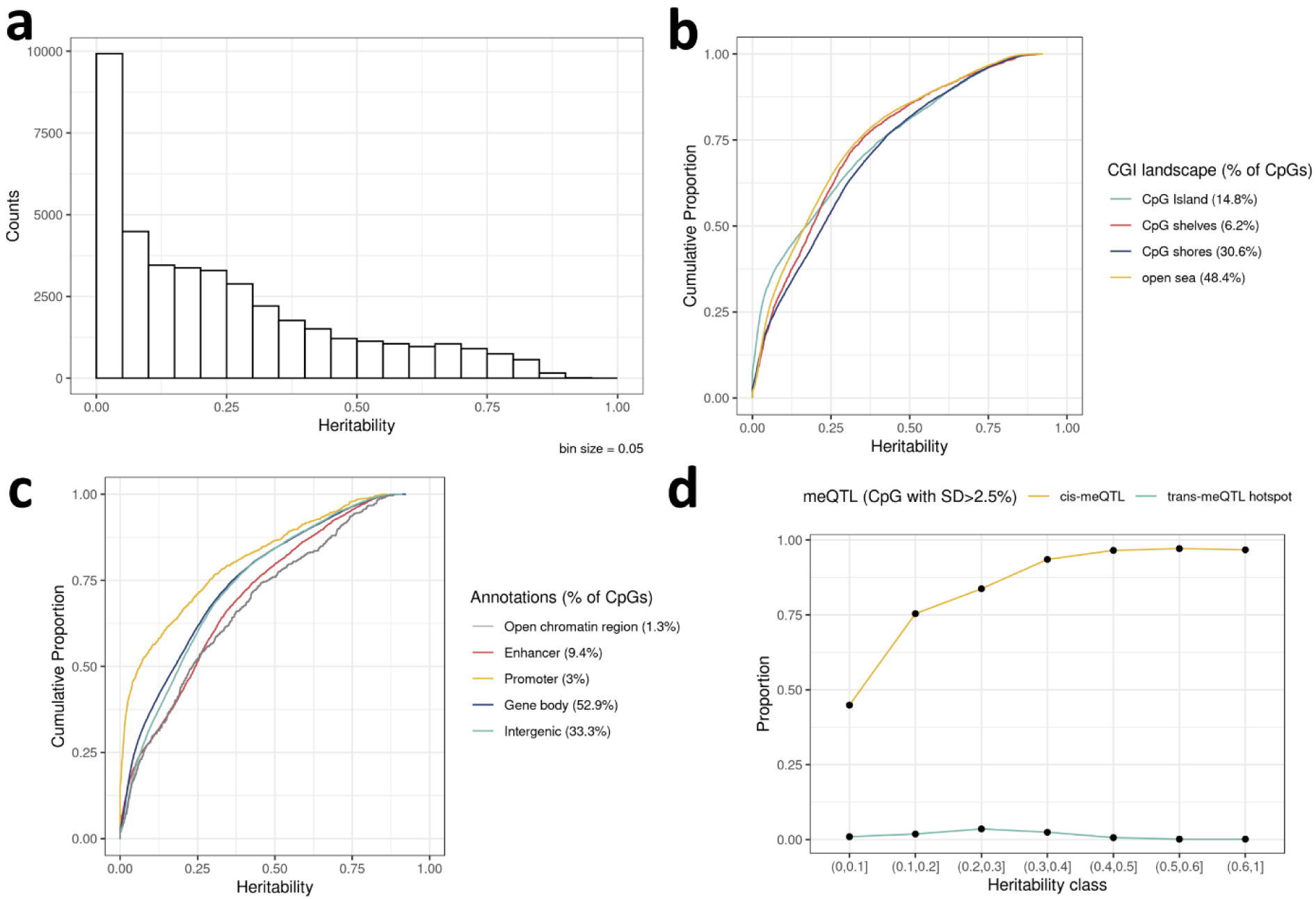
Heritability of the blood methylome. (a) Heritability distribution for 40,676 CpGs. (b) Cumulative distribution of heritability depending on the CpG island landscape (n=40,676 CpGs). (c) Cumulative distribution of heritability depending on the regulatory feature (n=40,676 CpGs). (d) Proportion of *cis*-meQTL and *trans*-meQTL hotspots for different heritability classes among the 28,806 CpGs with SD > 2.5%.

### Local CpG-SNP associations

Among the 28,806 CpG sites with SD > 2.5%, 23,068 (80.1%) were associated with a *cis*-meQTL at a *P*-value threshold of 2.06 × 10^-5^ (FDR < 0.05). Typically, sentinel *cis*-SNPs (those with the lowest *P-values*) were located close to their corresponding CpG sites (median of the absolute distance = 28.6 kb; Fig. 3a). Of note, a decay in association strength with increasing genomic distance was observed (Additional file 2: Fig. S1). Overall, compared with CpG sites not associated with *cis*-meQTLs, *cis*-CpGs showed modest over- or under-representation for different genomic annotations (Fig. 3b). In particular, they were enriched in enhancer (OR = 1.30, 95% CI= [1.17−1.45]). Enrichment and depletion patterns observed for the corresponding sentinel *cis*-SNPs were also moderate (Fig. 3b), with OR exceeding 1.5 only observed for promoters and CpG islands compared to a background set of SNPs of equal size and with similar MAF and CpG-distance distributions (for promoters: OR = 1.87, 95% CI= [1.42−1.48]; for CpG islands: OR = 1.51, 95% CI= [1.34−1.71]).

**Figure 3.**
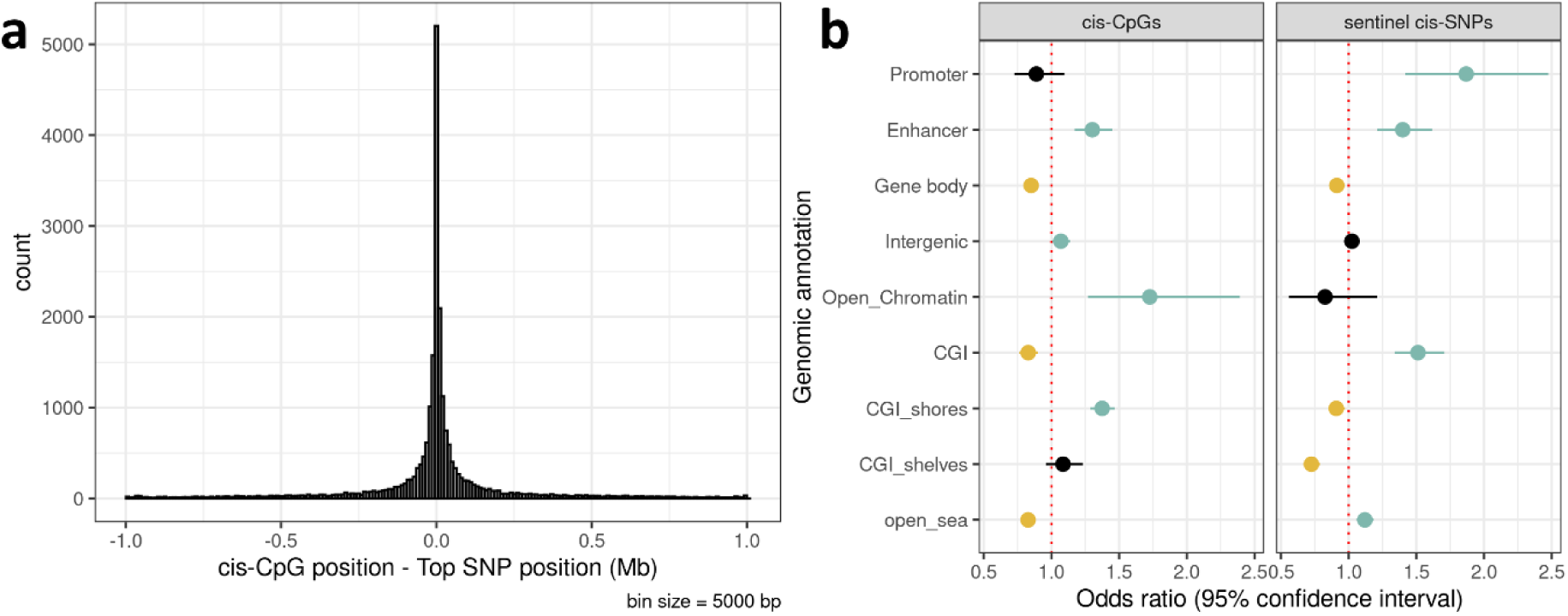
*cis*-meQTL mapping. (a) Distribution of the distance between *cis*-CpGs and their sentinel SNPs, in Mb. (b) Odds ratio and 95% confidence intervals of *cis*-CpGs and their corresponding sentinel *cis*-SNPs for several genomic annotations. OR are colored yellow when below 1, green when above 1, and black when not significantly different from 1.

### *Trans*-meQTL hotspots are associated with chromatin and transcription features

In this study, analyses of long-range associations were restricted to *trans*-meQTL hotspots, defined as genomic regions associated with multiple CpG sites (at least 34 independent *trans*-CpGs (>500 kbp apart); corresponding to the top 0.5% of SNPs; Fig. 4a). We allowed a permissive threshold for the initial *trans-*meQTL discovery step (P < 1×10^-6^). Only regions showing multiple co-localized associations were then further investigated, thereby limiting the selection of false positive associations. A total of 31 hotspots (denoted as H1 to H31) were identified, targeting 2,154 independent *trans*-CpGs (from 34 to 315 independent *trans*-CpGs per hotspot). The proportion of independent *trans*-CpGs represented on average 80.1% of the total number of *trans*-CpGs associated with the hotspot. Compared with the null expectation of 50%, the majority of hotspots (29 out of 31, 93.5%) exhibited a significant bias in the direction of effect across their associated *trans*-CpGs (*P* < 0.05, binomial test on independent *trans*-CpGs; Table 1). In other words, the polymorphism at the hotspot tended to primarily increase or decrease methylation levels of the associated *trans*-CpGs. Notably, for 21 hotspots (67.7%), the genes closest to the hotspots’ lead SNP have previously been associated with hematological traits in human GWAS [17]. Among these, associations were most frequently observed for platelet count (9/21; 41.8%), erythrocyte count (8/21; 38.0%), and eosinophil count (7/21; 33.3%) (Additional file 1: Table S2). We further investigated the genomic location of *tran*s-CpGs associated with each hotspot relative to several chromatin features, presented in Fig. 4b. Several hotspots exhibited strong enrichment patterns, which are detailed in the subsequent sections.

**Figure 4.**
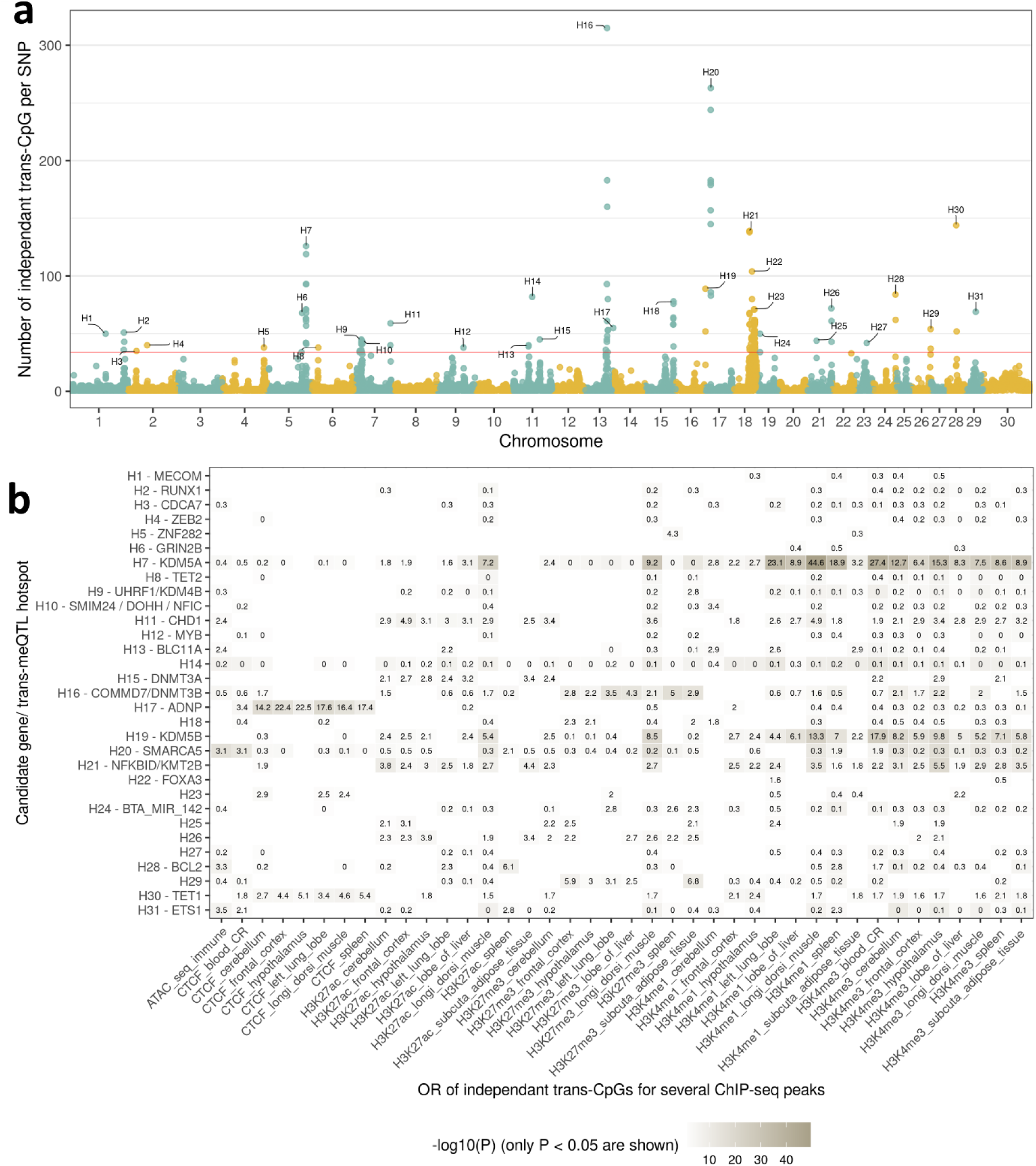
*Trans*-CpGs hotspots. (a) Manhattan plot of the number of independent *trans-*CpG with significant associations per SNP. Candidate genes are displayed for sentinel SNPs. Horizontal red line represents 34 independant *trans*-CpGs. (b) Enrichment of independent *trans*-CpGs for ATAC-seq peaks in immune cells, several histone marks and CTCF, for each hotspot. Odds Ratios (OR) are values inside boxes. Only OR with *P*-values < 0.05 are shown.

**Table 1.** ***Trans*-meQTL hotspots.**

| Hotspot | BT<br>A | Pos (50K) | lead PC1 SNP<br>(seq) | N ind (N) | Prop<br>"+" ind | TF | Dist<br>(kb) | P<br>(PWMEnrich) | P<br>(AME) | Candidate(s) |
| --- | --- | --- | --- | --- | --- | --- | --- | --- | --- | --- |
| H1 | 1 | 98125595 | 98122557 | 50 (58) | 0.42 | MECOM (MA0029.2) | 0 | 0.65 | 1 | MECOM |
| H2 | 1 | 147713803 | 147473912 | 51 (53) | 0.29 | RUNX1 (MA0002.3) | -229 | 5.5×10<sup>-3</sup> | 8.34×10<sup>-3</sup> | RUNX1 |
| H3 | 2 | 23461153 | 23340929 | 35 (39) | 1 | SP3 (MA0746.3) | -486 | 0.043 | 1 | CDCA7 |
| H4 | 2 | 52238741 | 52177770 | 40 (41) | 0.82 | ZEB2 (MA2686.1) | 0 | 0.10 | 1 | ZEB2 |
| H5 | 4 | 112324070 | 112307474 | 38 (44) | 0.29 | ZNF746 (MA2570.1) | 179 | 0.78 | 1 | ZNF282 |
| H6 | 5 | 96041071 | 96034274 | 68 (100) | 0 | / |  |  |  | GRIN2B |
| H7 | 5 | 106990076 | 107210779 | 126 (150) | 1 | TEAD4 (MA0809.3) | -376 | 0.07 | 1 | KDM5A |
| H8 | 6 | 20054035 | 20068167 | 38 (45) | 1 | / |  |  |  | TET2 |
| H9 | 7 | 19209398 | 19270709 | 45 (58) | 0.04 | RAX2 (MA0717.2) | 936 | 0.11 | 1 | UHRF1/KDM4B |
| H10 | 7 | 21196633 | 20437297 | 44 (48) | 0.07 | NFIC (MA1527.2) | 11 | 0.029 | 0.041 | SMIM24 / DOHH / NFIC |
| H11 | 7 | 98543164 | 98363092 | 59 (85) | 0.07 | / |  |  |  | CHD1 |
| H12 | 9 | 73264807 | 73260168 | 38 (44) | 0.97 | MYB (MA0100.4) | -2 | 0.65 | 0.99 | MYB |
| H13 | 11 | 43064231 | 43054225 | 40 (52) | 0.92 | BCL11A (MA2324.1) | 154 | 0.43 | 1 | BCL11A |
| H14 | 11 | 52758789 | 52652433 | 82 (83) | 0.02 | / |  |  |  | / |
| H15 | 11 | 73197270 | 74107406 | 45 (49) | 0 | / |  |  |  | DNMT3A |
| H16 | 13 | 62146317 | 62109421 | 315 (530) | 0 | PLAGL2 (MA1548.2) | -415 | 0.72 | 1 | COMMD7/DNMT3B |
| H17 | 13 | 78818064 | 78835159 | 55 (72) | 0.02 | SNAI1 (MA1558.2) | -793 | 0.22 | 1 | ADNP |
| H18 | 15 | 77233229 | 77184097 | 78 (100) | 0.17 | SPI1 (MA0080.7) | 17 | 0.49 | 1 | / |
| H19 | 16 | 79938997 | 79983101 | 89 (107) | 1 | LHX9 (MA0701.3) | -305 | 0.13 | 1 | KDM5B |
| H20 | 17 | 14265993 | 14238588 | 263 (390) | 0.83 | / |  |  |  | SMARCA5 |
| H21 | 18 | 46549874 | 46581441 | 139 (272) | 0.12 | ETV2 (MA0762.2) | -212 | 0.06 | 1 | NFKBID/KMT2B |
| H22 | 18 | 53388814 | 53398631 | 104 (124) | 0.22 | FOXA3 (MA1683.2) | -7 | 1.4×10<sup>-4</sup> | 1.0×10<sup>-4</sup> | FOXA3 |
| H23 | 18 | 58328337 | 58672622 | 71 (96) | 0.73 | / |  |  |  | / |
| H24 | 19 | 9302775 | 9302775 | 50 (60) | 0.18 | HSF5 (MA2676.1) | 90 | 0.91 | 1 | BTA_MIR_142 |
| H25 | 21 | 27654944 | 27692912 | 44 (51) | 0.7 | KLF13 (MA0657.2) | 0 | 1 | 1 | / |
| H26 | 21 | 67739884 | 67747776 | 72 (84) | 0.06 | / |  |  |  | / |
| H27 | 23 | 33848645 | 33919390 | 42 (43) | 0.07 | / |  |  |  | / |
| H28 | 24 | 59622599 | 61533118 | 84 (102) | 0.23 | / |  |  |  | BCL2 |
| H29 | 26 | 50757174 | 50836066 | 54 (96) | 0.02 | NKX6-2 (MA0675.2) | 177 | 0.94 | 1 | / |
| H30 | 28 | 23822036 | 24923672 | 144 (207) | 1 | ATOH7 (MA1468.1) | 868 | 0.10 | 1 | TET1 |
| H31 | 29 | 31048046 | 31025989 | 69 (81) | 0.52 | ETS1 (MA0098.4) | 788 | 0.26 | 1 | ETS1 |

Pos (50K): position of sentinel 50K variant associated with the largest number of independent *trans*-CpGs; lead PC1 SNP (seq): Position of the most significant sequence variant from the GWAS of the first coordinate of a PCA conducted on the *trans*-CpGs' hotspot; N ind (N): number of independent *trans*-CpGs, *i.e.*, *trans*-CpGs at least 500Kb apart (number of *trans*-CpGs); Prop "+" ind: proportion of independent *trans*-CpGs with a positive estimated $\beta$ effect; TF: transcription factor within 1 Mb of Pos (Seq) with the most significant PWMEnrich *P*-value; Dist (kb): Genomic distance in kb separating the TF from the most significant sequence variant; *P* (PWMEnrich): *P*-value corresponding to the TF motif enrichment in the independent *trans*-CpGs associated with the hotspot compared to the background sequences (independent *trans*-CpGs associated with all other hotspots), considering 100-bp sequences centered on the *trans*-CpGs obtained from PWMEnrich; *P* (AME): *P*-value corresponding to the TF motif enrichment in the independent *trans*-CpGs associated with the hotspot compared to the background sequences (independent *trans*-CpGs associated with all other hotspots), considering 100-bp sequences centered on the *trans*-CpGs obtained from AME; Candidate: candidate gene.

TFs and their binding sites can mediate long-range variant-CpG effects [10,13,14]. To investigate this mechanism in cattle, we performed motif enrichment analyses on the sequences surrounding *trans*-CpGs (100 bp centered on each CpG) for each hotspot, using *trans*-CpGs sequences associated with all other hotspots as the background set. For three hotspots, we observed both PWMEnrich and AME significant enrichment of *trans*-CpGs whose corresponding TF genes were located within 1 Mb of the sentinel SNP (*RUNX1*, *NFIC* and *FOXA3*; Table 1). The *RUNX1* gene was located 229 kb from the lead variant of hotspot H2. The *NFIC* and *FOXA3* genes were much closer to their respective hotspot lead variants, at 11 kb and 7 kb, respectively. Notably, *FOXA3* has previously been reported as a *trans*-meQTL hotspot in human blood [18].

### The COMMD7/DNMT3B locus

H16 was the *trans*-meQTL hotspot associated with the highest number of *trans*-CpGs (n = 530 with *P* < 1×10^-6^), encountered on chromosome 13, at 62.1Mb, with the estimated effect always in the same direction. Among its independent *trans*-CpGs, 37/315 were located on the X chromosome, a proportion significantly higher than observed for the other hotspots (OR = 10.1, *P* = 8.1 × 10^-18^, one-sided Fisher Exact Test). For this hotspot, several enrichments were observed in regulatory profiling datasets, including both ChIP-seq and CUT&RUN peaks, especially for H3K27me3 modification for several tissues. The lead SNP was located within *COMMD7*, and close to *DNMT3B*. *COMMD7* has been previously reported to be a *trans*-meQTL hotspot in human blood [13], linked to the basophile composition. *DNMT3B* is another compelling candidate, given its role as a *de novo* DNA methyltransferase. Its function is consistent with potential interactions between DNA methylation and H3K27me3-mediated chromatin repression [19] and the presence of multiple *trans*-CpGs targeting the X chromosome.

### The ADNP, SMARCA5 and TET1 loci

Several *trans*-meQTL hotspots showed enrichment of their *trans*-CpGs for CTCF binding sites in blood (Fig. 4b). CTCF is a key chromatin architectural protein involved in insulator activity, enhancer-promoter interactions, and higher-order genome organization, supporting its potential role in long-range epigenetic regulation [20,21]. H17 was associated with 72 *trans*-CpGs (55 independent *trans*-CpGs), with SNP 13:78835159C>A being the lead SNP for the sequence-level GWAS of PC1, located into the chromatin remodeler *ADNP*. The associated *trans*-CpGs showed an enrichment in CTCF-binding sites in multiples tissues including blood (OR = 3.4; *P* = 4.5 × 10^-5^). Previous work reported an overlap between CTCF- and ADNP-binding sites for Th2 cells [22] and ES cells [23]. Another hotspot, H20, composed of 390 *trans*-CpGs (263 independent *trans*-CpGs), was associated with a locus on chromosome 17 (lead SNP = 17:14238588:A>G) located in proximity to *SMARCA5* (Fig. 5a), with an OR of 3.1 for CTCF binding sites in blood (P = 1.8×10^-16^), consistent with the established role of *SMARCA5* in facilitating CTCF binding [24]. Functional enrichment analysis of H20-associated *trans*-CpGs revealed significant overrepresentation within genes involved in leukocyte differentiation (enriched term 25/256; FDR adjusted *P* = 1.9 × 10^-6^), lymphocyte differentiation (enriched term 20/256; FDR adjusted *P* = 1.9 × 10^-6^), and additional biological processes related to hematopoiesis and immune system regulation (Fig. 5b and Additional file 1: Table S3). These findings are further supported by prior studies implicating *SMARCA5* in hematopoietic stem and progenitor cell (HSPC) development [25,26]. We found another locus on chromosome 28 (H30) for which *trans*-CpGs preferentially overlap with CTCF binding sites in blood (OR = 1.8; P = 0.007). The medium density SNP was located at 23.8 Mb and was associated with 207 *trans-*CpGs (144 independent *trans*-CpGs). The sequence-level GWAS of PC1 of these *trans*-CpGs revealed SNP 28:24923672:G>C as the sentinel SNP, localized within *TET1*, a gene coding for a methylcytosine dioxygenase. The associated *trans*-CpGs all had the same estimated directional effect, with the C alternative allele increasing methylation. Of note, interactions between *TET1* and *CTCF* binding sites have been reported [27,28].

**Figure 5.**
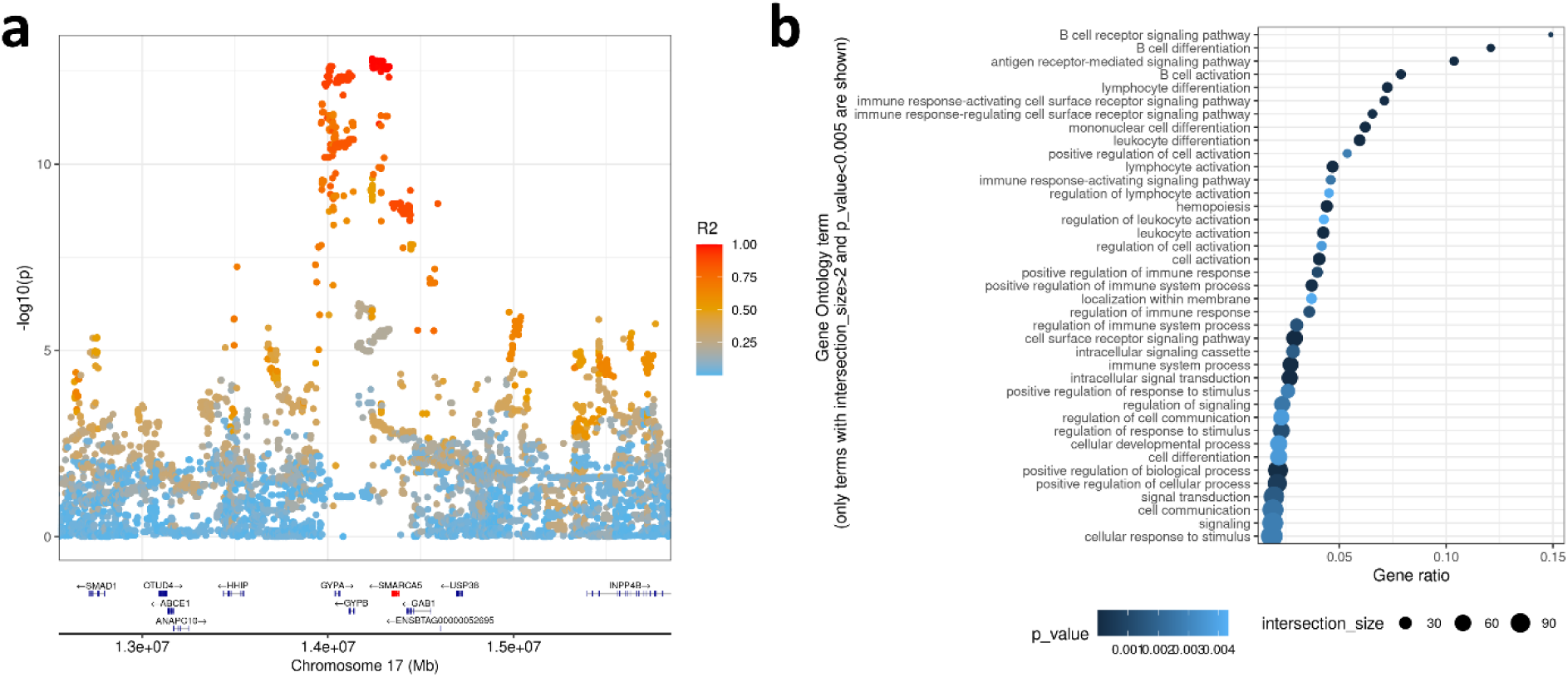
The H20/*SMARCA5 trans*-meQTL hotspot locus. (a) Regional Manhattan plot of the GWAS performed on the first principal component summarizing methylation variation across 390 *trans*-CpGs associated with the sentinel SNP 17:14238588:A>G (hotspot H20). *SMARCA5* is colored red (bottom). (b) Enrichment in Gene Ontology terms (“Biological Processes” only) for genes close to (<500 kb) *trans*-CpGs associated with H20. Only terms with an intersection size above 2 are shown.

### The *KDM4A* and *KDM5B* loci

The sentinel SNP 5:106990076A>G (50K level) of H7 was associated with 150 *trans*-CpGs (126 independent *trans*-CpGs, with the alternative allele always increasing methylation. At the sequence level, GWAS of PC1 of the 150 *trans-*CpGs highlighted a region containing *KDM5A* (Fig. 6b). *KDM5A* demethylates trimethylated and dimethylated H3K4 [29]. The associated independent *trans*-CpGs were located on multiple chromosomes (Fig. 6a) and were found to overlap with H3K4me3 marks more frequently than those associated with other hotspots in blood (OR = 27.4; *P* = 1.5 × 10^-41^; Fig. 6d). H3K4me3 is a chromatin mark typically enriched at active promoters and associated with transcriptional initiation, making it a relevant feature for widespread regulatory effects [30]. A second hotspot, H19 (sentinel 50K SNP 16:79938997A>G), was associated with 107 *trans*-CpGs (89 independent *trans*-CpGs) spread across multiple chromosomes (Fig. 6a), with the alternative allele always increasing methylation. Sequence level GWAS of PC1 of the associated *trans*-CpGs identify *KDM5B*, another histone demethylase of the KDM5 family, as a positional candidate gene (Fig. 6c). *KDM5B* is also involved in the methylation and demethylation of H3K4 [29]. As with the *KDM5A* locus, a higher proportion of independent *trans*-CpGs at this hotspot overlapped with narrow H3K4me3 peaks in blood than was observed for independent *trans*-CpGs associated with all other hotspots (OR = 17.8; *P* = 3.5 × 10^-28^; Fig. 6d). Genes closest to H19-associated *trans*-CpGs were overrepresented for multiple Gene Ontology terms including catalytic and transferase activities (Additional file 1: Table S3).

**Figure 6.**
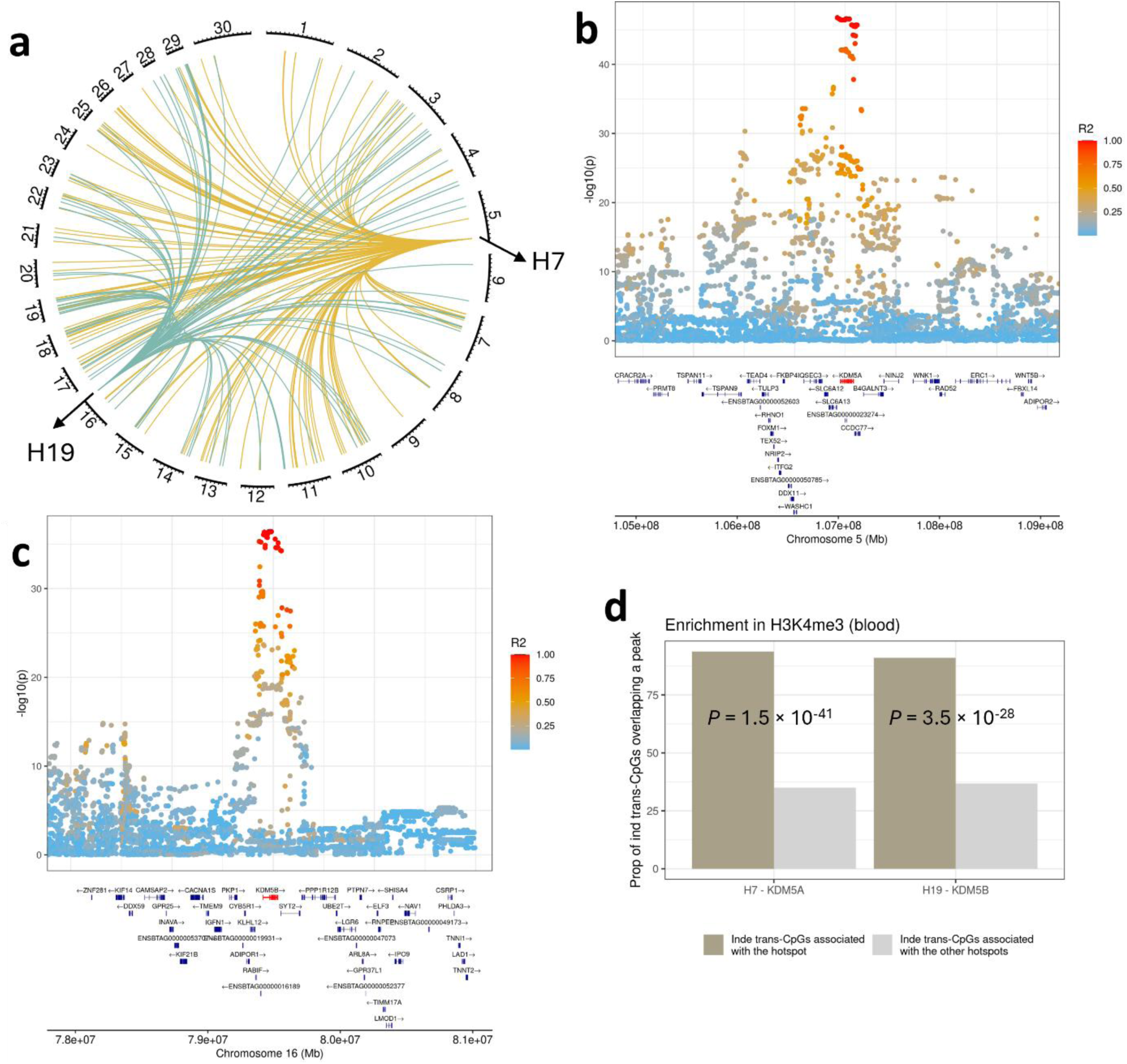
The *KDM5A* and *KDM5B trans*-meQTL hotspots loci. (a) Circos plot representing genomic locations of the *trans*-CpGs associated with the trans-meQTLs hotspot H7 (Chr5:106,990,076; 50K sentinel SNP; yellow) and H19 (Chr16:79,938,997; green). (b) Regional Manhattan plot of the GWAS performed on the first principal component summarizing methylation variation across 150 *trans*-CpGs associated with the sentinel SNP 106990076A>G. *KDM5A* is colored red (bottom). (c) Regional Manhattan plot of the GWAS performed on the first principal component summarizing methylation variation across 107 *trans*-CpGs associated with the sentinel SNP 16:79938997A>G. *KDM5B* is colored red (bottom). (d) Percent of independent *trans*-CpGs (at least 500Kb apart) associated with *KDM5A*/*KDM5B* hotspots overlapping with H3K4me3 narrow peaks measured in bovine blood. Background: independent *trans*-CpGs linked to other hotspots (grey).

### No evidence of colocalization between *trans*-meQTL hotspots and QTLs for common dairy traits

To investigate their potential effects on complex traits, we performed colocalization analyses in genomic regions surrounding *trans*-meQTL hotspots, integrating GWAS data for seven traits: milk somatic cell score, clinical mastitis, milk yield, milk fat yield, milk protein yield, heifer fertility, and cow fertility. Two hotspots (H5 and H21) showed a posterior probability of a shared causal variant with heifer fertility above 80%, while for both, the heifer fertility QTL did not reach the significant threshold (*P* > 5 × 10^-8^; Additional file 2: Fig. S2).

## Discussion

Leveraging data from 4,457 Holstein cows, this study represents, to our knowledge, the largest investigation of the genetic determinism of DNA methylation conducted in cattle and, more broadly, in livestock species. Overall, DNA methylation heritability estimates were moderate and most genetic effects on methylation operated over short genomic distances. In addition, our two-step strategy for mapping *trans*-meQTL hotspots enabled the precise identification of positional candidate genes, while the integration of functional genomic annotations provided strong support for several hotspot regions and their potential regulatory mechanisms.

To date, only one study has estimated the heritability of DNA methylation in cattle. Conducted in Holstein bull sperm using reduced representation bisulfite sequencing and restricted to variable CpG sites (SD > 5%), that study reported an average heritability estimate of 0.26 [31], which is comparable to our findings. Notably, these estimates are higher than those reported in humans, where blood DNA methylation heritability estimates typically range from ∼0.10 to 0.20 [11,12,14]. Such differences may partly reflect disparities in the genomic representation and design of epigenotyping arrays. Consistent with previous human studies, we observed lower heritability in promoter associated CpGs and higher heritability in enhancer-associated CpGs regions, potentially reflecting differences in DNA methylation levels or regulatory dynamics between these genomic regions.

Gunasekara et al. [32] previously identified correlated regions of systemic interindividual variation (CoRSIVs) in humans and demonstrated that these regions are under strong genetic control. The existence of CoRSIVs has also been reported in cattle and shown to be associated with genetic variation [16], in agreement with our results showing higher heritability estimates for CpGs located within CorSIVs. We also identified a substantially higher proportion of CpGs associated with *cis*-meQTLs (80.1% of variable CpGs) than has been reported in human whole blood studies [14,18], where only approximately one third of CpGs are influenced by nearby genetic variants. As previously mentioned, methodological differences between studies, including differences in CpG selection criteria and epigenotyping arrays, may contribute, at least in part, to these discrepancies.

A major finding of this study is the identification of several *trans*-meQTL hotspots associated with chromatin regulation and transcriptional processes. Previous studies have shown that *trans* effects on DNA methylation can arise through transcription factors and variations in their binding sites [12,14,18]. We observed similar patterns for several hotspots, including H22 involving *FOXA3*, a TF previously identified as a *trans*-meQTL hotspot in human blood [18] and *RUNX1* (hotspot H2) which has been implicated in *trans*-regulatory methylation networks [18]. Histone modifications constitute another plausible mechanism associating genetic variants to CpG methylation levels [33]. In rats, Rintisch et al. [34] demonstrated that histone QTLs can influence gene expression. Our results further suggest that histone modifiers may also affect CpG methylation at multiple distal genomic regions, as illustrated by hotspots H7 and H19. In addition, several hotspots mapped to regions containing genes playing a key role in DNA methylation or genes known to interact with them (*CDCA7* (H3), *TET2* (H8), *UHRF1* (H9), *DNMT3A* (H15), *DNMT3B* (H16), *BTA-MIR142* (H24), *TET1* (H30)). Among these, *BTA-MIR142*, a microRNA-encoding gene, emerged as a compelling candidate underlying *trans*-meQTL hotspot H24. *BTA-MIR142* is involved in hematopoietic and immune cell regulatory pathways and has been shown to target *TET2* [35], a key enzyme mediating active DNA demethylation. We therefore hypothesize that the *trans*-methylation effects associated with H24 may arise from a *cis*-regulatory variant affecting *MIR142* expression, which could in turn modulate *TET2* activity in *trans*, thereby influencing methylation levels at numerous CpGs distributed across the genome. Notably, the lead SNP (Chr19:9302775) co-localizes with a CTCF-binding site identified in blood using CUT&RUN data (Additional file 1: Table S4). Given the established role of CTCF in shaping chromatin architecture and regulating gene expression, local chromatin organization may influence *MIR142* expression. However, we cannot exclude the possibility that this hotspot partly reflects interindividual differences in blood cell composition rather than a direct regulatory effect on DNA methylation. More generally, although integrative analyses enabled the prioritization of candidate genes and biological mechanisms, experimental validation will be required to confirm these hypotheses.

When molecular QTLs are analyzed in bulk tissues, particularly *trans*-hotspots, detected associations may reflect loci influencing cellular composition rather than direct molecular regulation [3]. Although we adjusted for three major cell types, this correction remains incomplete given the numerous blood cell populations present in whole blood. Therefore, genes involved in hematopoiesis or immune-cell abundance represent plausible candidates underlying meQTL hotspots detected in blood. Although evidence in cattle remains limited, many of the positional candidate genes identified in this study have previously been associated with hematological traits in humans [17]. These hotspots therefore represent promising candidates for the genetic regulation of blood-related phenotypes in cattle, although additional studies will be necessary to validate these associations and clarify the underlying biological mechanisms.

### Limitations

This study has several limitations. First, the two-step approach used to identify long-range associations focused on *trans*-meQTL hotspots, therefore masking more specific long-range SNP-CpG regulatory relationships. In addition, our colocalization analyses were restricted to *trans*-meQTL hotspots and selected phenotypic traits of interest. The absence of transcriptomic data for the studied cohort also precluded the identification of CpG sites associated with variations in gene expression and limited our ability to investigate causal regulatory cascades linking genetic variation, DNA methylation, and gene expression. Although public resources such as cattleGTEx are available, these datasets are heterogeneous and meta-analytic in nature and do not provide matched methylation and transcriptomic measurements from the same blood samples, thereby limiting their integration with our analyses.

## Conclusions

In summary, this study provides the first large-scale integrative characterization of the genetic architecture underlying blood DNA methylation in a livestock species, building on recently developed functional annotation resources in cattle. Our findings demonstrate that DNA methylation is substantially influenced by genetic variation, with regulatory mechanisms that largely mirror those observed in humans, including strong *cis* regulation and the presence of biologically meaningful *trans*-meQTL hotspots. These results contribute to a better understanding of epigenetic regulation in cattle and provide a valuable framework for future studies investigating the molecular basis of complex traits. More broadly, they emphasize the importance of integrating complementary “omics” layers, particularly transcriptomic and chromatin-related data, to disentangle causal regulatory mechanisms and clarify the phenotypic consequences of epigenetic variation.

## Methods

### Animal cohort description

The study is based on 4,457 female Holsteins, with an average age of 3.9 years (SD = 1.6 years) and born between 2008 and 2023 (median = 2020). The animals were sampled from 247 herds, with an average of 16.2 animals sampled per herd. The biological material used for genotyping was ear cartilage or jugular blood, and for epigenotyping, blood collected from the tail vein. The experimental protocols were approved by the national ethics committee under the references APAFIS #40364-2023011317188842 v4, APAFIS #43155-2023042513537531 v3, APAFIS #40530-2023012613547159 v3 and APAFIS #40472-2023012010502425 v3.

### Genotypes

All cows were genotyped using medium density SNP arrays, including the BovineSNP50 Beadchip (Illumina, San Diego, CA, USA) and the EuroGMD Beadchip (Illumina, San Diego, CA, USA, https://www.eurogenomics.com/actualites/the-eurog-md:-a-unique-genotyping-microarray-for-cattle-.html). Imputation to whole-genome sequence level was done in two steps, as described in Sanchez et al. [36]: (1) from medium-density genotypes to 777 K SNPs using the BovineHD BeadChip (Illumina, San Diego, CA, USA) and FImpute [37], and (2) from 777 K SNPs to whole-genome sequence variants using Minimac [38]. Only variants with a Minimac imputation accuracy coefficient of determination (r^2^) greater than 0.3 and a Minor Allele Frequency (MAF) above 1% were kept. The ARS-UCD1.2 bovine genome sequence was used as a reference [39].

### Methylation values

DNA extraction, bisulfite conversion and epigenotyping were subcontracted to GD Biotech (France). All cows were epigenotyped using the RUMIGEN EpiChip array (Illumina, San Diego, CA, USA). Fluorescence intensities associated with each probe were reported within a raw intensity files (.idat) and were preprocessed using the “SeSAMe” R package [40] through the “openSesame()” function, with the prep argument set to “QCDPB”. Preprocessed methylation signals were subsequently converted into *β*-values for downstream analyses.

The proportions of the three major leukocyte cell types — neutrophils, lymphocytes, and monocytes — were estimated for each sample using a reference-based deconvolution approach implemented in the “EpiDISH” R package [41]. Reference DNA methylation profiles for each cell type were established in a previous study (Costes et al., personal communication; reference DI-RV-24-0038, INRAE Intellectual Property and Valorization Committee).

DNA methylation levels obtained with the RUMIGEN EpiChip were subjected to quality control and pre-treatment. Methylation marks with more than 20% missing values were discarded. The *β*-values of the remaining CpG sites were transformed using a rank-based inverse normalization and subsequently analyzed using a linear mixed model (“lme4” package in R). The model included age in days as a covariate, blood cell counts (proportions of neutrophils, monocytes, and lymphocytes) as covariates, and slide ID and slide position as random effects. The residuals from this model were retained and considered as phenotypes in subsequent analyses.

### Heritability estimation

For each CpG site, the proportion of genetic variance was estimated using REML method implemented in GCTA (v1.93.2) [42], integrating the genomic relationship matrix (GRM) computed from medium density autosomal SNPs. Heritability was defined as: 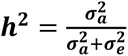 with 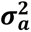 being the genetic additive variance and 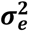 being the residual variance.

### Methylation QTL mapping

*Cis-* and *trans*-meQTL mapping were performed for 28,806 CpG sites with SD*_β_*_-value_ > 2.5% and SNPs with a MAF above 1%. SNP-CpG associations were tested using a mixed linear model implemented in the GCTA software (option “--mlma”; v1.93.2) [42], including the effect of the tested SNP and a polygenic random effect based on the medium density GRM. The resulting GWAS *P*-values were corrected for genomic inflation using the λ parameter estimated at the medium-density level. Fig. 1 displays the meQTL mapping design.

*Cis*-meQTL mapping was done within a 2 Mb window centered on each CpG site using sequence-level genotypes. To account for multiple testing, we controlled the false positive rate (FDR) at 5%. To achieve this, we first generated a distribution of *P*-values based on the null hypothesis by permuting animal identifiers (equivalent to permuting phenotypes) while retaining the GRM to preserve the population structure. Five permutations were performed, and for each permutation the lowest *P*-value per CpG site was retained. The significance threshold (T) was then adjusted so that, on average across the permutations: 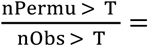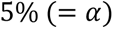, where nPermu corresponds to the number of CpG-SNP associations in the permuted set, and nObs to the number of CpG-SNP associations observed in the non-permuted set, considering the lowest *P*-value per CpG site in each case.

*Trans*-meQTL mapping was done in two steps due to the computational load intensity associated with using a GRM to correct for population structure:

1. *trans*-meQTLs mapping was first conducted at medium density using medium density genotypes (including the X chromosome), applying a permissive discovery threshold of *P* < 1×10^-6^. *Trans*-associations located on the same chromosome within 20 Mb (SNP-CpG distance) were removed, as they likely corresponded to long-range *cis* associations.
2. *trans*-meQTL hotposts were defined as the top 0.5% SNPs associated with the highest number of independent *trans*-CpGs (see next paragraph), corresponding to at least 34 independent *trans*-CpGs. For each hotspot, we performed a principal component analysis (PCA) on the *trans*-associated CpGs, and then conducted a local-sequence-level GWAS on the first principal component (PC1) within a 4 Mb region centered on the sentinel SNP (SNP associated with the largest number of *trans*-CpGs at medium density level) (Fig. 1). The rationale is that PC1 captures the major shared genetic component underlying variation across the associated *trans*-CpGs. Individuals with more than 20% missing values across these *trans*-CpGs were removed from the PCA. Remaining missing values were imputed by the mean.

We distinguished between independent and dependent *trans*-associations. For each SNP, the strongest association with a *trans*-CpG (the sentinel CpG) was designated as independent. All other *trans*-CpGs associated with the same SNP and located within 500 kbp of the sentinel CpG were classified as dependent. This procedure was applied recursively, selecting the next strongest remaining association as a new sentinel, until no *trans*-CpG remained.

### Low alignment quality CpGs removal

*Trans*-CpGs associated to numerous *trans*-meQTL hotspots were visually investigated for potential alignment issues (regions of high or low coverage compared to neighboring regions). Briefly, for 2 sequenced Holstein bulls, we used SAMtools (v1.21) [43] to extract a 1 Mb region centered on the CpG(s) and visualized the “.bam” files using IGV (v2.19.7) [44]. CpG sites located in such low-quality regions were excluded from subsequent analyses.

### Annotations and enrichments

#### Curated databases

Annotation of CpGs and SNP were performed using Ensembl (v109) [45] for gene location, UCSC genome browser for CpG islands (CGI) coordinates (“cpgIslandExt” table) [46]. To further characterize the CGI landscape, two additional regions around the CpG islands were defined: (1) CGI shore regions, located 0-2 kb from CpG islands and (2) CGI shelf regions, located 2-4 kb regions from CpG islands. CpGs and SNPs outside these regions were assigned as belonging to the “open sea” category. Additionnaly, we used regulatory profiling datasets (see “Regulatory profiling datasets” section) and we retrieved ATAC-seq peaks from Yuan et al. [5]. In their study, Yuan et al. [5] applied non-negative matrix factorization (NMF) to decompose chromatin accessibility profiles across tissues into latent components representing tissue-specific regulatory profiles. In the present study, we kept only peaks with positive weights in the “Immune system” component, therefore corresponding to regulatory elements showing accessibility patterns enriched in immune-related cell types.

#### Regulatory profiling datasets

Publicly available ChIP-seq datasets from bovine samples annotated in Kern et al. [4]. In that study, genome-wide functional annotation was performed across eight adult bovine tissues, including liver, lung, skeletal muscle, spleen, subcutaneous adipose, cerebellum, brain cortex, and hypothalamus, generating comprehensive maps of regulatory elements in cattle. The epigenomic profiling included five ChIP-seq assays per tissue targeting CTCF and four histone modifications (H3K4me3, H3K27ac, H3K4me1, and H3K27me3), enabling the identification of promoters, enhancers, and repressive chromatin states. More details about the method utilized are available in Kern et al. [4].

Because blood-derived cells were not utilized to build the regulatory map, we additionally generated CTCF and H3K4me3 profiles from a whole blood sample collected from a Holstein cow and stored at −80 °C. CTCF binding sites and H3K4me3 profiles in whole blood were generated using the CUT&RUN (Cleavage Under Targets and Release Using Nuclease) protocol. An *in-house* optimized CUT&RUN protocol was applied using approximately ∼100,000 nuclei per reaction. Buffers from the CUT&RUN Assay Kit (Cell Signaling Technology; #86652) were used in combination with magnetic beads (Polysciences; #86057) and pAG-MNase (Curie Institute; PR-E-003). CTCF antibodies were obtained from Diagenode (C15410210). Library preparation was performed using the SimpleChIP® ChIP-seq DNA Library Prep Kit for Illumina® (#56795) and SimpleChIP® ChIP-seq Multiplex Oligos for Illumina® – Dual Index Primers (#4753). Libraries were sequenced on an Illumina NovaSeq platform (100 bp paired end). CUT&RUN raw sequencing data were analyzed using the nf-core/cutandrun pipeline (v3.2.2; https://github.com/nf-core/cutandrun).

#### Enrichments of *cis*-CpGs and their *cis*-meQTLs

Enrichment of *cis*-CpGs and their sentinel *cis*-meQTL SNPs (lowest *P*-value) for genomic annotations were assessed using Fisher’s exact test. For *cis*-CpGs, the background set consisted of all the CpGs retained for the meQTL mapping without a *cis*-meQTL. For the *cis*-meQTLs top SNPs, the background set consisted of an equal number of SNPs randomly sampled from all variants tested in the *cis*-meQTL analysis. Background SNPs were matched to the foreground set based on both MAF (10 bins spanning 0–100% in 10% increments) and SNP-CpG distance (10 bins: [−1 Mb, −500 kb), [−500 kb, −100 kb), [−100 kb, −50 kb), [−50 kb, −10 kb), [−10 kb, 0 kb), [0 kb, 10 kb), [10 kb, 50 kb), [50 kb, 100 kb), [100 kb, 500 kb), and [500 kb, 1 Mb]).

#### Enrichments of *trans*-CpGs targeted by *trans*-meQTL hotspots

*Trans*-CpGs targeted by *trans*-meQTL hotspots were subjected to enrichment analysis for regulatory elements. Enrichments were performed considering independent *trans*-CpGs associated with the hotspot as the foreground set and independent *trans*-CpGs associated with all other hotspots as the background set. Enrichment for the histone marks, CTCF and ATAC-seq peaks were assessed using Fisher’s exact test. Enrichment analyses for known transcription factors (TF) binding motif sequences were performed using the “PWMEnrich” R package [47] and AME (MEME suite; v5.5.4) [48]. For each hotspot, we tested whether the 100 bp sequences centered on the corresponding *trans*-CpGs were enriched for motifs of TFs encoded within 1 Mb of the hotspot lead SNP. The background set consisted of the centered 100 bp sequences of *trans*-CpGs associated with all other hotspots. For PWMEnrich, we computed a log-normal background distribution and used the threshold-free affinity score. For AME, the average odds scoring method was applied, with *P*-values obtained from Fisher’s exact test. The TF list and position frequency matrices were retrieved from JASPAR 2026 [49], restricting the analysis to non-redundant vertebrate TFs. *P*-values below 0.05 were considered significant. Additionally, genes located near *trans*-CpGs associated with meQTL hotspots were tested for functional enrichment using the “gprofiler2” R package [50]. Data sources search was restricted to the Gene Ontology and KEGG databases. For each *trans*-CpG, the closest gene within 500 kb was retained using the closest function from BEDTools [51]. Only enrichments with FDR corrected *P*-values below 0.05 were considered significant.

### Colocalization analysis

We explored the colocalization between *trans*-meQTLs hotspots and QTLs for seven traits related to health, production and fertility: milk somatic cell score, clinical mastitis, milk yield, milk fat yield, milk protein yield, heifer fertility, and cow fertility. GWAS for traits of interest were performed on daughter yield deviations of 8,792 to 10,066 Holstein bulls, depending on the trait [52]. Colocalization analyses were conducted using the “coloc” R package [53]. For each significant QTL (*P* < 5 × 10^-8^) located within a 4 Mb window centered on the trans-meQTL hotspot, we retained only analyses for which PP.H3.abf (posterior probability of distinct causal variants) was below 0.5 and PP.H4.abf (posterior probability of a shared causal variant) was above 0.8.

## Declarations

### Ethics approval and consent to participate

Not applicable.

### Consent for publication

Not applicable.

### Availability of data and materials

Restrictions apply to the availability of the original data supporting the findings of this study due to third party ownership. Genotypes data of the 4,457 female Holsteins were produced for the purpose of bovine selection and belong to French breeding organizations, which have given INRAE permission to use them for research purposes, but are not publicly available. The curated datasets were obtained from Kern et al. [4] (ChIP-seq dataset), Yuan et al. [5] (ATAC-seq). GWAS results for the seven traits (milk somatic cell score, clinical mastitis, milk yield, milk fat yield, milk protein yield, heifer fertility, and cow fertility) originates from Charles et *al.* [52] (https://doi.org/10.1038/s41467-025-58970-5) and can be downloaded from the Recherche Data Gouv database (https://doi.org/10.57745/UO9T9O). FASTQ files of sequenced Holstein animals used to detect alignment problems of CpGs and SNP are available in the European Nucleotide Archive (ENA) at EMBL-EBI (https://www.ebi.ac.uk/ena/browser/view/) under accession number PRJEB64022 (sample ID: SAMEA120832690) and PRJEB64023 (sample IDs: SAMEA114111987, SAMEA114111990). Additional files 1 to 4 will be made available upon acceptance of the article.

### Authors’ contributions

C.F., D.B. and M.-P.S. conceived the study design. S.F., C.P. H.K., D.B., M.-P.S. supervised the project. C.F. performed most data analysis, with contribution from V.C. and G.C.M.M. C.F. V.C., F.B., G.C.M.M., M.B., H.K., S.F., C.L.D and M.-P.S. contributed to data interpretation. M.J. contributed to optimise the CUT&RUN protocol for blood samples. C.F. wrote the initial version of the paper. All authors read and approved the final manuscript.

## Acknowledgements

We acknowledge the breeding companies that provided biological samples and access to cows’ genotypes. Imputations were carried out with sequences from the “1000 Bull Genomes” project. We thank G. Potier, L. Le Berre, C. Richard and V. Gélin for blood collection, and F. Ali, A. Asset, A. Bonnet, A. Chaulot-Talmon, M. C. Deloche and A. Raja Ravi Shankar for their support with preparing and handling the samples used in this study.

